# Cortical projection neurons exhibit early maturation defects and selective degeneration in a mouse model of spinal muscular atrophy

**DOI:** 10.64898/2026.10.09.758065

**Authors:** Roberta Schellino, Federica La Rocca, Giulia Iezzi, Anna Caretto, Marina Boido, Alessandro Vercelli

## Abstract

Spinal muscular atrophy (SMA) is usually considered a lower motor neuron disease; however, growing evidence from both experimental models and clinical studies indicates that SMN deficiency affects the whole CNS, beyond the spinal and brainstem motor system. Here, we investigated the temporal progression and selective vulnerability of cortical projection neurons in the SMNΔ7 mouse model at early (P5) and late (P11) symptomatic stages. Combining subtype-specific immunohistochemistry, DiI tracing associated with morphometric reconstruction, and dendritic spine analysis, we identified stage- and subtype-dependent cortical alterations. At P11, corticospinal and infragranular callosal projection neurons were selectively reduced in number and showed marked dendritic remodeling and altered excitatory synaptic organization, whereas corticothalamic neurons and supragranular SATB2-positive populations were comparatively preserved. Notably, cortical abnormalities were already detectable at P5, before overt neuronal loss: indeed, corticospinal neurons exhibited reduced soma size and dendritic complexity, while dendritic spine density remained preserved, but spine morphology was shifted toward immature phenotypes. These early maturation defects persisted during disease progression and preceded the reduction in spine density observed at P11. Together, our findings identify cortical projection neurons as selectively vulnerable to constitutive SMN deficiency and suggest that impaired neuronal and synaptic maturation represents an early feature of cortical pathology preceding subtype-specific neurodegeneration. These results expand the pathophysiological spectrum of SMA beyond lower motor neurons and highlight early brain alterations as a potential component of disease progression.

**Graphical Abstract:** *Cortical projection neuron vulnerability and early maturation defects in SMA:* Schematic summary of cortical alterations in SMNΔ7 mice. Early structural and synaptic maturation defects are already present in corticospinal neurons at P5 despite preserved projection neuron populations, whereas by P11 these abnormalities are accompanied by reduced spine density and selective degeneration of vulnerable layer V projection neurons (corticospinal and corticocortical). These findings suggest that postnatal developmental defects in neuronal and synaptic maturation represent an early marker of cortical vulnerability to SMN deficiency, preceding the subtype-specific neurodegeneration observed at later disease stages. Created with BioRender.com.

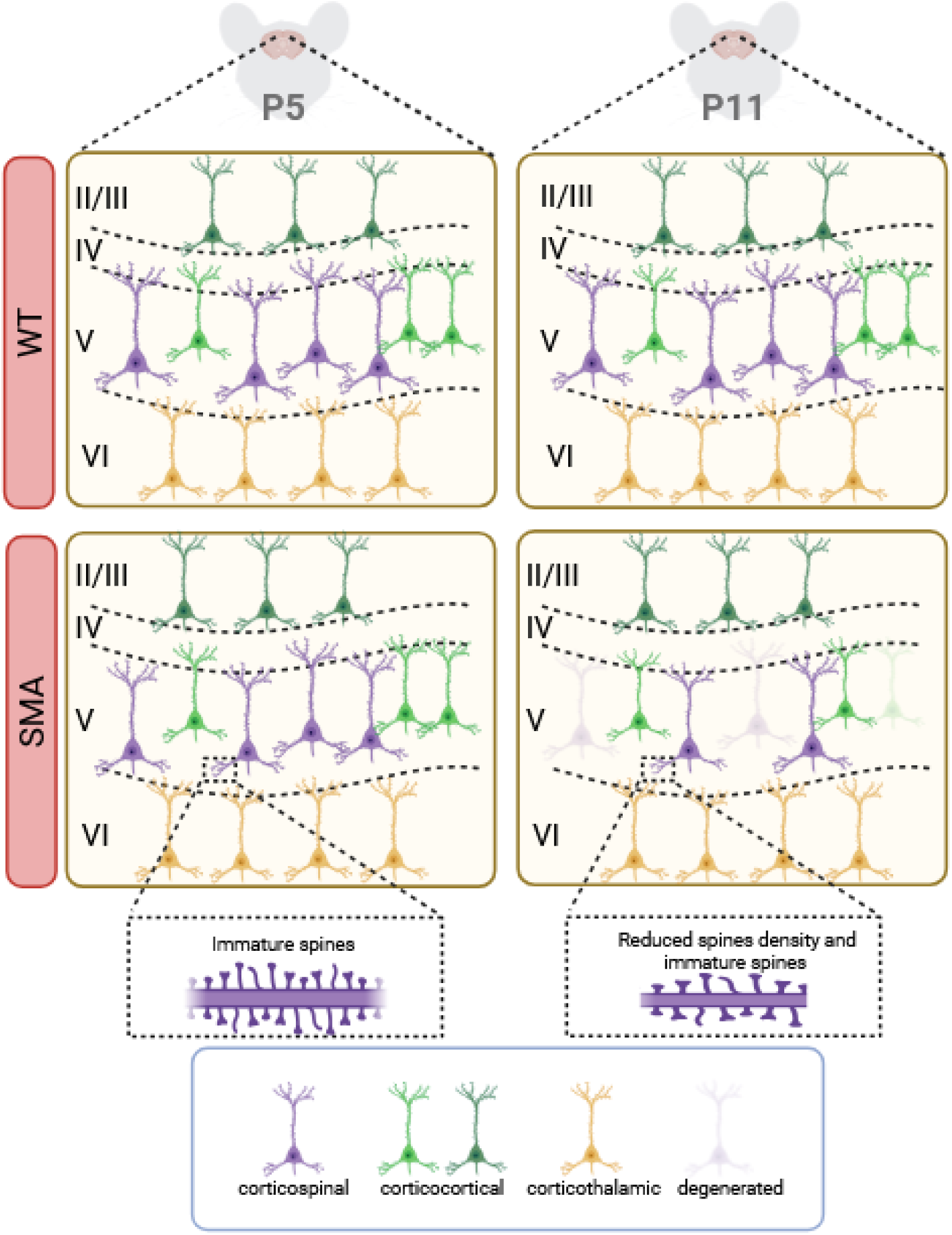

## Introduction

Spinal muscular atrophy (SMA) is an autosomal recessive neuromuscular disorder caused by homozygous deletion or mutation of the *Survival Motor Neuron 1* (*SMN1*) gene on chromosome 5q, resulting in markedly reduced levels of functional SMN protein. The nearly identical paralog SMN2 produces only limited amounts of full-length SMN because of predominant exon 7 skipping, and its copy number is a major modifier of disease severity, contributing to the broad clinical spectrum of SMA (Lefebvre et al., 1995, 1997; Lorson et al., 1999, 2010; Verhaart et al., 2017).

Historically, SMA has been considered a selective lower motor neuron disorder, characterized by progressive degeneration and loss of spinal and brainstem motor neurons, skeletal muscle denervation, progressive atrophy, and weakness (Lorson et al., 1999). Extensive evidence from human tissue and animal models, such as the SMNΔ7 mouse, has established spinal motor neuron loss and selective vulnerability of specific motor neuron pools as key features of disease pathology (Burghes and Beattie, 2009; d’Errico et al., 2013; Martinez et al., 2012; Schellino et al., 2018). However, the classical motor unit-centered view has progressively expanded. Indeed, SMN protein is ubiquitously expressed throughout peripheral tissues and the central nervous system (CNS) (Hamilton and Gillingwater, 2012; Nash et al., 2016), and beyond its canonical role in small nuclear ribonucleoprotein assembly and RNA processing, participates in cytoskeletal dynamics, axonal transport, growth cone function, and synaptic maintenance (Fallini et al., 2012; Thomsen et al., 2021; Shi et al., 2024). Thus, SMN deficiency may affect multiple neuronal populations and interfere with neuronal development and maintenance beyond the lower motor system.

Although neuropathological observations in humans are relatively limited, particularly to severe SMA forms, degenerative signs have been reported in several supraspinal regions, including the cerebral cortex, thalamic nuclei, brainstem and cerebellum (Harding at al., 2015; Mendonça et al., 2019; Maeda et al., 2019; Thomsen et al., 2021). Neuroimaging further support CNS involvement in SMA (Querin et al., 2018; Mugisha et al., 2023; Groulx-Boivin et al., 2024). Such findings suggest altered brain development or remodeling processes and challenge the traditional view that SMA spares supraspinal structures.

Experimental models have provided further evidence for brain involvement. In SMNΔ7 mice, abnormalities have been reported in both the motor cortex and cerebellum, including loss and functional impairment of selected neuronal populations and altered circuit organization (d’Errico et al., 2013; Tharaneetharan et al., 2020; Cottam et al., 2023). More recently, alterations in cortical inhibitory circuitry have also been identified (Menduti et al., 2026), while in vitro primary SMA cortical neurons display reduced viability and morphological abnormalities under basal conditions (Rasà et al., 2025), further supporting the intrinsic vulnerability of cortical neuronal populations to SMN deficiency. Complementary evidence from SMA I patient-derived cerebral organoids identified early abnormalities in neuronal activity (Faravelli et al., 2025), further indicating that early brain dysfunction is an intrinsic component of SMA pathogenesis.

Despite growing recognition of brain involvement, the extent and nature of cortical neuronal alterations in SMA are still not comprehensively characterized. In particular, little is known about the differential vulnerability of different cortical projection neuron populations. Previous studies have reported loss of layer V pyramidal neurons in SMNΔ7 mice (d’Errico et al., 2013) and abnormalities of Betz cells and corticospinal pathways in human SMA tissue (Araki et al., 2003), but whether distinct cortical projection neuron subtypes undergo early structural and synaptic alterations before overt degeneration remains unclear.

Here, we systematically characterize distinct populations of cortical projection neurons in the SMNΔ7 mouse model at early- and late-symptomatic stages. By combining quantitative, morphometric, and synapse analyses, we investigate the temporal emergence and subtype specificity of cortical alterations, with particular emphasis on neuronal morphology, dendritic organization, and spine maturation. This approach aims to define early features of cortical vulnerability to constitutive SMN deficiency and their relationship with the selective degeneration observed during disease progression and may provide new insights for therapeutic strategies extending beyond lower motor neuron rescue to include the broader neural networks affected by SMN deficiency.

## Materials and Methods

### Animals

SMNΔ7+/+; SMN2+/+; Smn+/− breeder pairs (stock no. 005025; The Jackson Laboratory, Bar Harbor, ME, USA) were used to generate the experimental cohorts. Intercrosses produced Smn−/− mice, which were used as a model of severe SMA, and Smn+/+ littermates serving as wild-type (WT) controls. Newborn pups were identified by tail snip at postnatal day 0 (P0), and genotyping was performed by PCR analysis following the supplier’s protocol. Animals were maintained with the mother until sacrifice at P5 or P11 (early and late disease stage, respectively) and housed under standard conditions (12 h light/12 h dark cycle) with unrestricted access to food and water. Both male and female pups were included in the study. All procedures were designed to minimize animal use and discomfort. Experiments were conducted in compliance with current European regulations (Directive 2010/63/EU) and Italian law and policies governing animal research (D.L. N.26, 04/03/2014) and were approved by the Italian Ministry of Health (authorization #160/2020-PR) and by the Ethical Committee of the University of Turin. A total of 79 mice (39 WT and 40 SMA) were used. All the analyses were performed blinded to genotype.

### Tissue preparation

For immunofluorescence, immunohistochemistry, and DiI (1,1’-dioctadecyl-3,3,3’,3’-tetramethylindocarbocyanine perchlorate) tracing experiments, WT and SMA mice at P5 and P11 were deeply anesthetized with isoflurane and transcardially perfused with 0.1 M phosphate buffer (PB, pH 7.4), followed by cold 4% paraformaldehyde (PFA, Carl Roth, CAS No. 30525-89-4) in 0.1 M PB (pH 7.4). Brains were carefully removed from the skull and post-fixed in 4% PFA for 1h at 4°C. Tissues used for immunofluorescence and immunohistochemistry analysis were then cryoprotected overnight in 30% sucrose (Sigma-Aldrich, CAS No. 57-50-1) dissolved in 0.1 M PB until complete sinking. After cryoprotection, brains were embedded in cryostat medium (Killik, Bio-Optica, Milan, Italy) and frozen. Coronal brain sections were cut at the desired thickness using a cryostat (Leica). Free-floating sections were stored at −20°C in antifreeze solution (30% ethylene glycol, 30% glycerol in 0.1 M PB) until further processing. In parallel, following postfixation, tissues to be used for for DiI tracing were stored in 0.1M phosphate buffer saline (PBS) containing 0.02% sodium azide until further processing.

### Histochemistry and immunofluorescence

For Nissl staining, 40µm-thick brain sections were mounted onto 2% gelatin-coated slides and air-dried overnight. Sections were rehydrated in distilled water and stained with 0.1% Cresyl violet acetate (Sigma-Aldrich). Following staining, sections were dehydrated through ascending ethanol solutions, cleared in xylene, and coverslipped with Eukitt mounting medium (Bio-Optica).

For immunofluorescence analysis, free-floating sections (40 µm) were first rinsed in PBS to remove the antifreeze solution and subsequently permeabilized for 30 minutes in PBS containing 0.3% Triton X-100. Following permeabilization, non-specific binding sites were blocked for 30 minutes in PBS containing 0.3% Triton X-100 and 10% normal serum. Sections were then incubated for 24 h at 4°C with primary antibodies diluted in 0.1 M PBS (pH 7.4) containing 0.3% Triton X-100 and 2% appropriate normal serum. The following primary antibodies were used: CTIP2 (AB18465; rat; Abcam, Cambridge, UK); SATB2 [SATBA4B10] (AB51502; mouse; Abcam, Cambridge, UK); TBR1 (AB31940; rabbit; Abcam, Cambridge, UK); MAP2 (188006; chicken; Synaptic System, Gottingen, Germany); cleaved Caspase3 (CASP3; rabbit; Cell Signaling Technology, Danvers, MA, USA); vesicular glutamate transporter-1 (vGlut1; 135 302; Synaptic System, Gottingen, Germany); NeuN (MAB377; mouse; Millipore, Burlington, Massachusetts, USA). After primary antibody incubation, sections were washed in PBS and subsequently incubated for 2 hours with the corresponding fluorophore-conjugated secondary antibodies (Jackson ImmunoResearch Laboratories, West Grove, PA, USA). Nuclei were counterstained with DAPI (4′,6-diamidino-2-phenylindole) for 3 min at room temperature. After being washed in PBS, sections were then mounted onto 4% gelatin-coated slides, coverslipped with the antifade mounting medium Mowiol, and examined using a Zeiss LSM 980 confocal microscope.

### DiI tracing and morphometric measurements

Fixed brains were cut sagittally into slices (2-3 mm thick) using a mouse brain matrix. Crystal of the lipophilic tracer Cy3-DiI (Thermo Fisher Scientific, Cat. No. D-282) were dissolved into 99% Ethanol (UN1170, Sial) to obtain a concentrated dye preparation, following the manufacturer’s recommendations. A small amount (≃4µl) of the DiI preparation was aspirated into a microsyringe and locally delivered into selected white matter tracts. The amount of tracer deposited at each site was adjusted empirically until a clearly visible, spatially restricted dye deposit was obtained. DiI was locally applied to selected white matter tracts at defined rostrocaudal levels, identified according to anatomical landmarks of the P5 mouse brain. Tracer deposits were placed in the cerebral peduncle at the midbrain level to retrogradely label corticospinal/corticofugal projection neurons, in the corpus callosum at an intermediate rostrocaudal level to label callosal projection neurons, and in the dorsal thalamic region to label corticothalamic projection neurons. After dye application, sections were then immersed in 4% PFA and incubated at 37°C for 3–4 weeks to allow diffusion of the tracer along neuronal membranes. Brain samples were subsequently included in 4% agar (Lab Box, Cot. No. AGAR-D1L-500), then sectioned at 100 µm on a Leica VT1200S vibratome (Leica Microsystems) (speed 0.35 mm/s; frequency ∼70 Hz). Individual sections were mounted onto 4% gelatin-coated slides and analyzed immediately after cutting without coverslipping, in order to minimize lateral dye diffusion and preserve signal localization (Vercelli et al., 2000). Mounting media were avoided, as they reduced DiI fluorescence under our imaging conditions and could promote dye diffusion or dissolution, thereby compromising signal localization.

For morphometric reconstruction, only DiI-labeled pyramidal neurons were selected according to laminar position and stringent morphological criteria. Corticospinal neurons were analyzed in layer V, corticothalamic neurons in layer VI, and callosal projection neurons within infragranular layers (mainly upper layer V). Only neurons exhibiting unequivocal pyramidal morphology (triangular soma, clearly identifiable apical dendrite, and well-resolved dendritic arborization) and sufficient isolation were included. Cells were excluded when DiI-labeled processes from neighboring neurons overlapped with the cell of interest, potentially confounding reconstruction. Soma size, total neurite length, and dendritic branching were reconstructed using Neurolucida software (MBF Bioscience, Williston, VT), and quantitative parameters were extracted with NeuroExplorer (MBF Bioscience). For basal dendrites, both total and mean length were quantified: total length was calculated as the summed length of the entire dendritic arbor, including all branches arising from successive branch points, whereas mean length represented the average length of individual dendritic segments between branch points. Neurite complexity was quantified by the software as a dimensionless index reflecting branching structure (complexity index). For each projection neuron subtype analyzed, DiI-labeled cells were obtained from at least three independent animals per genotype (N ≥ 3).

### Golgi-Cox staining

Golgi-Cox staining was performed according to a modified protocol adapted from Zaqout and Kaindl (2016). Briefly, fresh brains were collected from WT and SMA mice at P5 and P11 immediately after cervical dislocation and briefly rinsed in distilled water. Whole brains from P5 and hemisected brains from P11 mice were processed. Tissue samples were immersed in freshly prepared Golgi–Cox impregnation solution (10 ml per sample) composed of 5% potassium dichromate (K₂Cr₂O₇), 5% mercuric chloride (HgCl₂), and 5% potassium chromate (K₂CrO₄) in distilled water. Brains were kept at room temperature in the dark. After 24 h, the impregnation solution was replaced with freshly prepared solution, and tissues were maintained under the same conditions for 7–10 days (depending on tissue-size). Following impregnation, samples were transferred to a cryoprotectant solution consisting of 30% sucrose, 1% polyvinylpyrrolidone (PVP-10), and 30% ethylene glycol in phosphate buffer. After 24 h, tissues were placed in fresh cryoprotectant solution and stored at 4°C in the dark for an additional 4–7 days. Then, brains were embedded in 4% agarose and sectioned coronally at 100 µm using a Leica VT1200 S vibratome (speed 0.35 mm/s; frequency ∼70 Hz). The vibratome chamber was filled with cryoprotectant solution to preserve tissue integrity during cutting. Sections were mounted directly onto 4% gelatin-coated slides, gently dried to remove excess solution, and air-dried in the dark for 2–3 days.

For visualization of the Golgi reaction, sections were sequentially rinsed in distilled water, incubated in 50% ethanol for 5min, treated with diluted ammonia solution (3:1 in distilled water; 8min) to enhance signal development, and then, after being rinsed twice in distilled water, fixed for 10min in 5% sodium thiosulfate under dark conditions. Sections were subsequently dehydrated through graded ethanol solutions (70%, 95%, 100%; 6 min each), cleared in xylene, and coverslipped using Eukitt mounting medium (Bio-Optica). Slides were dried horizontally in the dark for 48 h prior to imaging. Golgi-Cox-impregnated neurons were analyzed from at least four independent animals per genotype (N ≥ 4).

### Dendritic spine measurement and counting

Morphological analyses were conducted using Neurolucida software (MBF Bioscience) interfaced with a Nikon Eclipse E600 light microscope equipped with a 100× oil-immersion objective. For each animal, five Golgi-Cox-impregnated pyramidal neurons located in layer V of the sensorimotor cortex were selected based on complete impregnation and absence of truncation or interferences from overlapping processes.

Dendritic spine density and morphology were evaluated on basal dendrites and on the primary apical dendrite of each neuron. Spine quantification was performed within standardized segments proximal to the soma, defined as 10 µm from the soma for basal dendrites and 20 µm for the apical dendrite. Spine density was expressed as the number of spines per micrometer of dendritic length. In addition, the distance between the soma and the first identifiable spine was measured along each analyzed dendrite. Spines were morphologically classified according to established criteria (Pchitskaya and Bezprozvanny, 2020) into four categories representing increasing levels of maturation: filopodia, stubby, thin, and mushroom.

### Quantitative Analysis

#### Nissl-based quantification of layer V projection neurons

Quantitative analyses of projection neuron density in cortical layer V were performed on Nissl-stained coronal sections at P5 and P11. For each animal, three anatomically matched sections encompassing motor cortex (mCTX) and the adjacent somatosensory cortex (sCTX) were selected and acquired under brightfield illumination using a E800 Nikon Eclipse light microscope equipped with a 20× objective. Sections were matched to the corresponding plates of the Allen Mouse Brain Atlas (M1 and S1; https://mouse.brain-map.org/static/atlas), and cortical regions were identified according to atlas-defined anatomical landmarks. Layer V was identified based on cytoarchitectonic criteria. Images were calibrated in Fiji/ImageJ (NIH) using the acquisition scale (µm/pixel), enabling area measurements in mm². Each image was converted to 8-bit grayscale and Nissl-positive somata were segmented using intensity thresholding (manual adjustment to optimally capture neuronal somata), followed by threshold application to obtain a binary mask. When required, binary masks were refined using morphological operations and separation of touching objects. Automated soma counting was performed within the image, applying a minimum size filter (size 15 to infinity for P11 brains; size 10 to infinity for P5 brains) to selectively include large somata consistent with pyramidal projection neurons, while excluding smaller Nissl-positive profiles likely corresponding to interneurons or glial cells. Objects intersecting image borders were excluded. Cell density was calculated as the number of counted neurons divided by the image area (cells/mm²). For each animal, densities from the three sections were averaged to obtain a single mean value for statistical analysis.

Genotype-related differences (or lack thereof) observed with ImageJ were independently confirmed by blinded manual quantification by Neurolucida software (MBF Bioscience), tracing layer V boundaries and counting large pyramidal somata with clearly identifiable cell bodies to compute densities (cells/mm²).

#### Immunofluorescence-based quantification of projection neuron populations

Quantification of specific projection neuron subtypes was performed on confocal images (40× magnification,1 μm *z*-step size, acquisition speed 100 Hz, format 1024 × 1024 pixels) of immunofluorescent stained sections of P5 and P11 brains. Marker-positive nuclei were manually counted using ImageJ/FIJI (Cell Counter plugin). The analyzed markers included CTIP2 (corticofugal neurons), SATB2 (callosal/intracortical neurons), and TBR1 (corticothalamic neurons), in combination with DAPI nuclear staining. MAP2 immunostaining was used when necessary to assist in the identification of projection neuron somata and to exclude non-neuronal elements. For each animal, four representative coronal sections including the mCTX and the adjacent sCTX were selected. From each section, three non-overlapping fields within the region of interest (layer V-VI or supragranular layers, depending on the analysis) were acquired under identical confocal settings across experimental groups. Cell density was expressed as the number of marker-positive nuclei per mm². Only nuclei with morphological characteristics consistent with projection neurons (larger DAPI-positive nuclei with euchromatic appearance) were included in the counting; small, densely stained DAPI-positive nuclei were excluded from the analysis. For each animal, values obtained from the three fields per section were averaged, and then the mean of the four sections was calculated to generate a single value per animal.

### Quantification of Caspase-3 expression

To assess activated Caspase-3 (CASP3) immunoreactivity as a potential indicator of apoptotic signaling in projection neuron populations, the proportion of CASP3-positive neurons was quantified among CTIP2+ and SATB2+ cells in P5 and P11 brains. Confocal images (40× magnification,1 μm *z*-step size, acquisition speed 100 Hz, format 1024 × 1024 pixels) were acquired from four representative coronal sections per animal, and three fields per section were collected. For each field, the number of CASP3-positive nuclei co-expressing CTIP2 or SATB2 was counted and normalized to the total number of CTIP2+ or SATB2+ neurons within the same field. Results were expressed as the percentage of CASP3+/CTIP2+ or CASP3+/SATB2+ cells relative to the total CTIP2+ or SATB2+ population. For each animal, values from all fields and sections were averaged to obtain a single mean percentage value.

### Quantification of vGlut1 puncta and signal intensity

To evaluate glutamatergic inputs onto cortical neurons, vesicular glutamate transporter 1 (vGlut1) immunostaining was analyzed. For each animal, four representative coronal sections were acquired using confocal microscopy; from each section, z-stack images were obtained (63× magnification, 1 μm z-step size, acquisition speed 100 Hz, 1024 × 1024 pixel resolution). Z-stack acquisition allowed inclusion of only those NeuN-positive neurons fully contained within the optical volume and ensured visualization of all vGlut1-positive puncta contacting the selected cell. For each section, 10 NeuN+ neurons with clearly defined soma contours and without overlapping with neighboring cells were manually selected. The soma contour was manually traced, and vGlut1-positive puncta located within the traced area or along its perimeter (indicative of putative synaptic contacts) were manually counted using ImageJ software. The mean number of vGlut1 puncta per neuron was calculated for each animal.

In addition, for each confocal image, overall vGlut1 signal intensity was quantified using ImageJ. Briefly, after converting vGlut1 staining images in black and white and applying an automatic Gaussian blur filter to reduce background noise, a local intensity threshold was set using a defined radius around each pixel. The positive fractioned area within the field of view (i.e., the percentage of vGlut1-positive pixels relative to the total image area) was calculated. This parameter was used as a measure of global glutamatergic signal density within layer V. For each animal, puncta counts and positive fractioned area values were averaged across sections and fields before statistical analysis.

### Statistical Analysis

All data are presented as mean ± standard error of the mean (SEM). Statistical analyses were carried out using GraphPad Prism (version 8.4; GraphPad Software, San Diego, CA, USA).

For histological/histochemical and morphometric datasets, normality of data distribution was assessed using the Kolmogorov–Smirnov and Shapiro–Wilk tests. Comparisons between two groups were then performed using an unpaired Student’s t-test for normally distributed data. These analyses were applied to pairwise comparisons. For comparisons involving multiple groups or factors, two-way ANOVA was used when data met parametric assumptions and was applied to analyze dendritic spine morphology distributions, considering genotype and spine type (or dendritic compartment) as independent factors, followed by appropriate post hoc tests.

## Results

### Layer V corticospinal and callosally-projecting, but not corticothalamic neurons, are reduced in the SMA cortex at the late symptomatic stage

To investigate the impact of SMN deficiency on cortical projection neurons, we first assessed whether these neuronal populations undergo neurodegenerative alterations at the late symptomatic stage of the disease. To this aim, we performed quantitative analyses of distinct cortical projection neuron populations in the SMNΔ7 cortex at P11. We confirmed the reduction in pyramidal neurons in SMA cortical layer V (as previously observed in d’Errico et al., 2013), compared to WT, by Nissl-staining (WT 1154 ± 56.21 cells/mm^2^; SMA 763.9 ± 62.69 cells/mm^2^; t-test, p=0.003**; n= 4 WT; 5 SMA; Figures 1A-A’), both in the motor (mCTX) and somatosensory (sCTX) areas (mCTX, WT 1205 ± 86.24 cells/mm^2^; SMA 802.3 ± 86.47; t-test, p=0.014*. sCTX, WT 1117 ± 66.56 cells/mm^2^; SMA 726.3 ± 60.52; t-test, p=0.034**; Figure 1A’’). Blinded manual quantification using Neurolucida software confirmed the differences in layer V neuronal density between WT and SMA mice (not shown).

**Figure 1.**
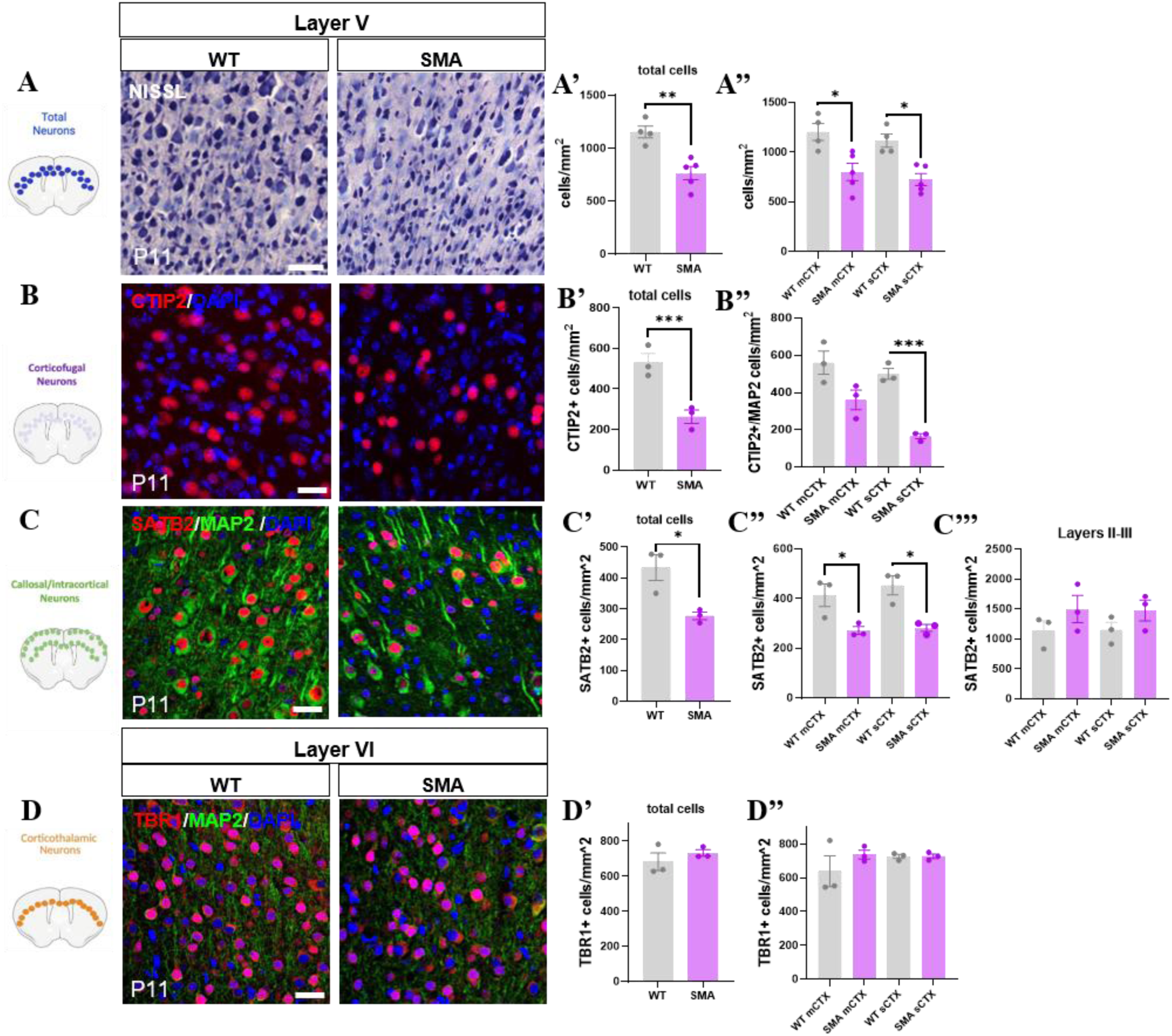
Reduction of cortical projection neuron populations in the SMA cortex at the late symptomatic stage (P11). (A) Representative Nissl-stained sections of motor cortex from WT and SMA mice at P11. (A’-A’’) Quantification of pyramidal neuron density in layer V reveals a significant reduction in SMA mice compared to WT in total and in both motor (mCTX) and somatosensory (sCTX) cortices. (B) Immunofluorescence analysis of CTIP2-positive corticofugal neurons in layer V. Quantification shows a significant decrease in CTIP2-positive neuron density in SMA cortex (B’), particularly in somatosensory areas (B’’). (C) Immunofluorescence and quantification of SATB2-positive callosal projection neurons reveal a significant reduction in layer V of SMA mice (C’) in both motor and somatosensory cortices (C’’). (C’’’) No significant differences are observed in SATB2-positive neurons in supragranular layers (II–III). (D) Analysis of TBR1-positive corticothalamic neurons in layer VI shows no significant differences between WT and SMA mice in total (D’) and in motor and somatosensory cortices (D’’). Data are presented as mean ± SEM. Scale bars: 50 µm.

Projection neuron subtype-specific immunofluorescence analyses demonstrated that this reduction is selective. Indeed, CTIP2-positive corticofugal neurons were significantly decreased in layer V of SMA mice, consistent with vulnerability of corticospinal projections (WT 531.3 ± 44.39 cells/mm^2^; SMA 263.5 ± 32.87 cells/mm^2^; t-test, p=0.0084**. n= 3WT; 3 SMA) (Figures 1B-B’). This reduction was particularly evident in the somatosensory areas (WT 501.6 ± 29.65 cells/mm^2^; SMA 166.0 ± 13.87 cells/mm^2^; t-test, p<0.001***), but a similar trend toward reduced CTIP2-positive neuron density was also observed in the mCTX of SMA mice (WT 561.0 ± 62.78 cells/mm^2^; SMA 361.0 ± 52.45 cells/mm^2^; t-test, p=0.07) (Figure 1B’’).

Similarly, SATB2-positive callosal projection neurons showed a marked reduction in density in SMA cortex layer V (WT 433.9 ± 41.80 cells/mm^2^; SMA 277.2 ± 12.33 cells/mm^2^; t-test, p=0.023*; n= 3 WT; 3 SMA. Figure 1C-C’), in both motor (WT 414.2 ± 45.59 cells/mm^2^; SMA 272.1 ± 15.39 cells/mm^2^; t-test, p=0.042*) and somatosensory (WT 453.5 ± 38.04 cells/mm^2^; SMA 282.4 ± 14.13 cells/mm^2^; t-test, p=0.013*) areas (Figure 1C’’). Interestingly, given that SATB2 is also expressed in neurons located in the supragranular layers, we additionally quantified SATB2-positive cells in layers II–III of both mCTX and sCTX. In contrast to the infragranular layer V findings, no significant differences (p>0.05) were detected in the density of supragranular SATB2-positive neurons between SMA and WT mice (Figure 1C’’’). This observation indicates that the degenerative process seems to selectively affect infragranular projection neurons, while supragranular corticocortical/callosal populations remain preserved at this stage of disease progression.

Consistent with this finding of selective vulnerability among projection neuron subtypes, TBR1-positive corticothalamic neurons located in layer VI did not exhibit significant differences between genotypes (WT 683.0 ± 49.79 cells/mm^2^; SMA 732.9 ± 18.30 cells/mm^2^; t-test, p>0.05; n= 3 WT; 3 SMA), in both mCTX and sCTX (p>0.05) (Figures 1D-D’’), indicating relative preservation of this projection neuron subtype.

To determine whether the reduction of corticofugal and callosally-projecting neurons at P11 reflects ongoing apoptotic processes, thus confirming neuronal death within specific projection neuron populations, we quantified Caspase-3 (CASP3) expression in defined cortical subtypes. At the late symptomatic stage, the proportion of CASP3-positive cells among CTIP2-positive corticofugal neurons was significantly increased in SMA mice compared to WT littermates in layer V (WT 0.14 ± 0.01% CASP3+/CTIP2+ cells; SMA 0.67 ± 0.08% CASP3+/CTIP2+ cells; t-test, p= 0.0025**; n= 3 WT; 3 SMA. Figures 2A-2B), in both mCTX and sCTX (Supplementary Fig. 1A). A similar increase was observed in SATB2-positive callosal neurons (WT 0.054 ± 0.02% CASP3+/SATB2+ cells; SMA 0.53 ± 0.12% CASP3+/SATB2+ cells; t-test, p= 0.017*; n= 3 WT; 3 SMA. Figures 2C-2D), in both motor and somatosensory areas (Supplementary Fig. 1B) indicating that both infragranular projection neuron populations exhibit enhanced apoptotic signaling in SMA animals.

**Figure 2.**
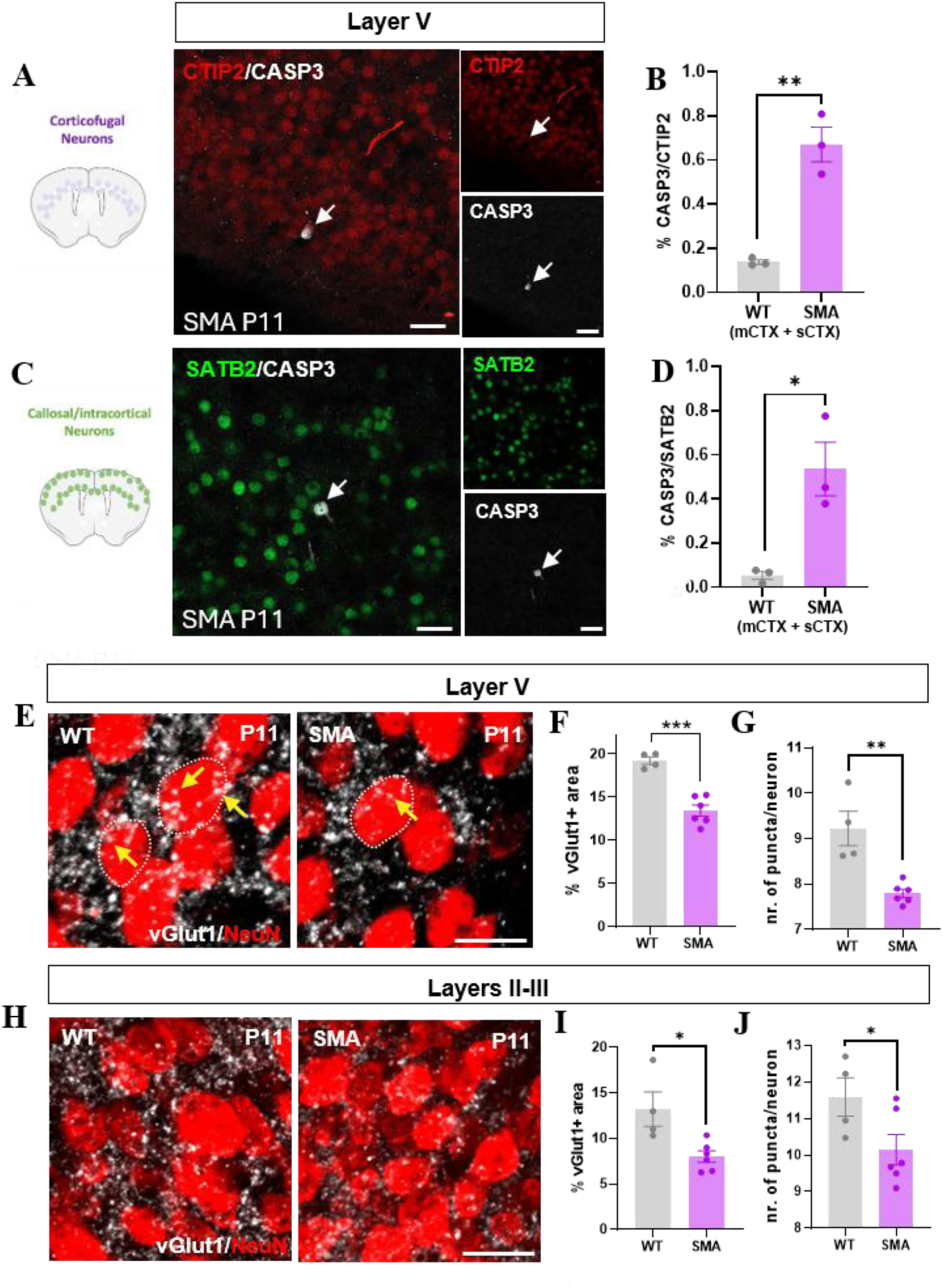
Increased apoptotic signaling and reduced excitatory synaptic input in cortical projection neurons of SMA mice at the late symptomatic stage (P11). (A-B) Representative images and quantification of CASP3 expression in CTIP2-positive corticofugal neurons in layer V of WT and SMA mice. The proportion of CASP3+/CTIP2+ cells is significantly increased in SMA cortex, indicating enhanced apoptotic signaling in corticofugal neurons. (C–D) Representative images and quantification of CASP3 expression in SATB2-positive callosal neurons, showing a significant increase in CASP3+/SATB2+ cells in SMA mice compared to WT, consistent with selective vulnerability of infragranular projection neuron populations. Representative images (E) and quantification of vGlut1-positive signal intensity (F) in layer V, revealing a significant reduction in SMA mice. (G) Quantification of vGlut1-positive puncta per NeuN-positive neuron in layer V, showing a decrease in SMA cortex. (H-I) Representative images and quantification of vGlut1-positive signal intensity in supragranular layers (II–III), demonstrating a significant reduction in SMA mice. (J) Quantification of vGlut1-positive puncta in supragranular layers, showing a decrease in SMA cortex. Data are presented as mean ± SEM. Scale bars: 50 µm (A, C); 20 µm (E, H).

To determine whether the degeneration of layer V projection neurons was accompanied by alterations in the local cortical circuitry, we next examined glutamatergic synaptic input in this layer. Quantitative analysis of vGlut1 immunoreactivity revealed a significant reduction in vGlut1-positive signal intensity within layer V of SMA mice compared to WT controls (WT 19.24 ± 0.43% vGlut1+ signal; SMA 13.44 ± 0.66% vGlut1+ signal; t-test, p= 0.0002***; n= 4 WT; 6 SMA. Figures 2E-2F). In addition to the overall decrease in glutamatergic signal, the number of vGlut1-positive puncta per NeuN-positive neuron was significantly reduced in SMA cortex (WT 9.22 ± 0.38 vGlut1+ puncta/ NeuN+ cell; SMA 7.80 ± 0.09 vGlut1+ puncta/ NeuN+; t-test, p= 0.0021**; n= 4 WT; 6 SMA. Figure 2G). This reduction was evident in both mCTX and sCTX (Supplementary Figures 1C-1D).

We next extended this analysis to supragranular cortical layers (II–III), where no significant changes in the density of SATB2-positive cortico-cortical projection neuron populations were observed. Quantitative analysis revealed a significant reduction in vGlut1-positive signal intensity in the supragranular layers of SMA mice compared to WT controls (WT 13.24 ± 1.88% vGlut1+ signal; SMA 8.02 ± 0.64% vGlut1+ signal; t-test, p= 0.015*; n=4 WT; 6 SMA. Figures 2H-I), in mCTX and sCTX (Supplementary Fig. 1E). Similarly, the number of vGlut1-positive puncta was also significantly decreased (WT 11.59 ± 0.52 vGlut1+ puncta/ NeuN+ cell; SMA 10.17 ± 0.41 vGlut1+ puncta/ NeuN+; t-test, p= 0.018*; n=4 WT; 6 SMA. Figure 2J), especially in sensory areas (Supplementary Fig. 1F). Overall, reduced vGlut1 immunoreactivity across cortical layers suggests an altered excitatory synaptic environment in the SMA cortex at the late symptomatic stage.

### Corticospinal and callosal, but not corticothalamic neurons show subtype-specific morphological alterations in the late symptomatic SMA cortex

To further characterize structural changes underlying the selective reduction of projection neuron populations at P11, we performed detailed morphometric analyses of DiI-labeled cortical neurons using Neurolucida reconstruction. Owing to the early postnatal age of the animals, DiI tracing could be reliably performed directly on fixed brain sections, as the relatively low degree of myelination and reduced tissue density at this developmental stage favor efficient lipophilic dye diffusion (Honig and Hume, 1989; Vercelli at al., 2000) (Figures 3A-C).

**Figure 3.**
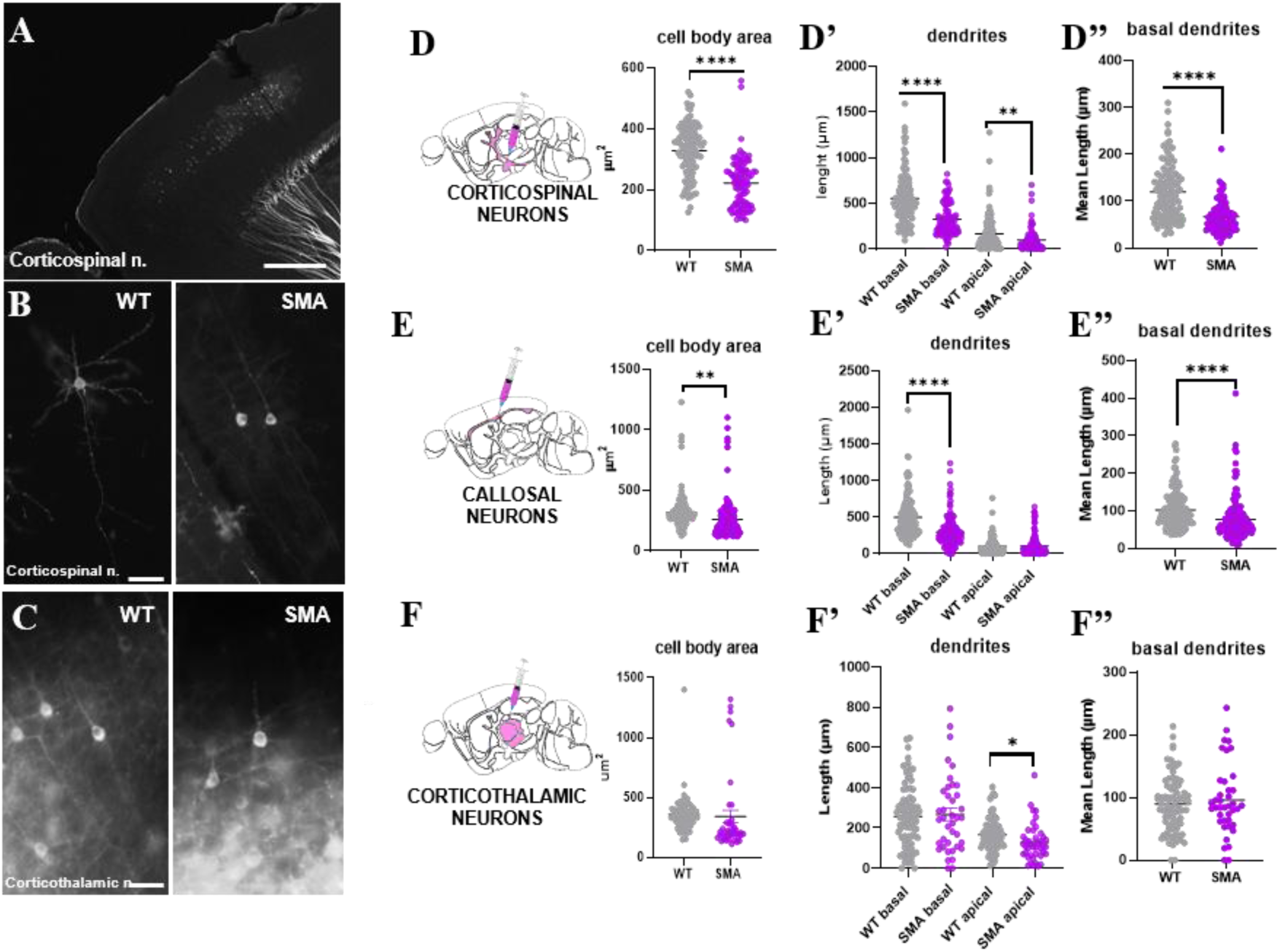
Morphological analysis of projection neuron cell types in the cortex reveals alterations in corticospinal and callosal, but not in corticothalamic, neurons at P11. (A-B) Representative fluorescence images of DiI-labeled corticospinal neurons in WT and SMA mice following DiI injection into the cerebral peduncle. (C) Representative fluorescence images of DiI-labeled corticothalamic neurons in WT and SMA mice following DiI injection into the thalamus. (D) Quantification of soma size reveals a significant reduction in SMA corticospinal neurons compared to WT. (D’–D’’) Quantification of dendritic architecture shows a marked reduction in total and mean basal dendrite length, as well as shortening of apical dendrites in SMA corticospinal neurons. (E) SMA callosal neurons exhibit reduced soma size and (E’-E’) shorter basal dendrites, while apical dendrite length is not significantly altered. (F-F’-F’’) No significant differences are observed between WT and SMA mice in corticothalamic neuron’s soma size and dendritic length. Data are presented as mean ± SEM. Data are presented as mean ± SEM. Scale bars: 500 µm (A); 50 µm (B-C).

At the late symptomatic stage, corticospinal neurons in the SMA cortex displayed evident morphological alterations compared to WT cells. Morphometric analysis of DiI-traced cells revealed a significant reduction in soma size (Table 1; n= 113 cells WT; 99 cells SMA. Figures 3D), accompanied by shorter basal and apical dendrites (Table 1; Figures 3D’, 3D’’). In addition, the dendritic arborization appeared less elaborate, with reduced branching and overall dendritic complexity, indicating structural remodeling of surviving corticofugal neurons compared to wild-type condition (Supplementary Fig. 2A-B).

**Table 1.** Morphological parameters of DiI-labelled corticospinal and corticothalamic neurons at P11.

| **Morphological Parameter** | **Wild Type**<br>**(Mean ± SEM)** | **SMA**<br>**(Mean ± SEM)** | **p-value**<br>**(t-test)** |
| --- | --- | --- | --- |
| ***Corticospinal neurons (Fig. 3 D- D'')*** |  |  |  |
| Soma Area ($\mu\text{m}^2$) | 328.4 ± 8.01 | 221.8 ± 8.26 | p<0.0001**** |
| Total basal dendrite length ($\mu\text{m}$) | 555.7 ± 26.53 | 327.4±17.46 | p<0.0001**** |
| Basal dendrites mean length ($\mu\text{m}$) | 119.6 ± 5.54 | 66.94 ± 3.23 | p<0.0001**** |
| Total apical dendrite length ($\mu\text{m}$) | 169.0 ± 17.95 | 100.4±12.10 | p=0.0024** |
| ***Callosally-projecting neurons (Fig. 3 E- E'')*** |  |  |  |
| Soma Area ($\mu\text{m}^2$) | 315.4 ± 11.14 | 259.4 ± 12.66 | p=0.001** |
| Total basal dendrite length ($\mu\text{m}$) | 498.0 ± 21.64 | 296.8 ± 16.21 | p<0.0001**** |
| Basal dendrites mean length ($\mu\text{m}$) | 102.5 ± 4.00 | 76.98 ± 4.51 $\mu\text{m}$ | p=0.0002*** |
| Apical dendrite length ($\mu\text{m}$) | 101.6± 8.434 | 104.1± 10.84 | p= 0.854 ns |
| ***Corticothalamic neurons (Fig. 3 F- F'')*** |  |  |  |
| Soma Area ($\mu\text{m}^2$) | 357.8± 16.19 | 342.0± 52.12 | p= 0.717 ns |
| Total basal dendrite length ($\mu\text{m}$) | 257.9± 17.14 | 266.8± 30.72 | p=0.786 ns |
| Basal dendrites mean length ($\mu\text{m}$) | 89.97±8.9 | 89.97±4.9 | p= 0.507 ns |
| Apical dendrite length ($\mu\text{m}$) | 166.3± 9.09 | 129 ± 14.30 | p=0.024* |

Similarly, callosally-projecting neurons in SMA mice exhibited marked morphological changes, suggesting compromised structural integrity. These neurons showed a reduced soma size (Table 1; n= 153 cells WT; 154 cells SMA. Figure 3E) and shorter basal dendrites (Table 1; Figures 3E-E’’), with a simplified dendritic architecture compared to WT controls (Supplementary Figures 2C-D). Apical dendrites of callosal neurons did not show significant differences in length between WT and SMA (p> 0.05). In contrast, corticothalamic neurons labeled by DiI injection into the thalamus display only minor morphological alterations. Quantitative analyses revealed minor differences between SMA and WT animals, with a modest but significant reduction in apical dendrite length, while soma size, basal dendritic length, and overall branching complexity were not significantly altered (n= 82 cells WT; 70 cells SMA; p> 0.05) (Table 1; Figure 3F-F’’ and Supplementary Fig. 2E-F). These findings are consistent with the preservation of TBR1-positive corticothalamic neuron density.

To further investigate the structural alterations of cortical projection neurons, dendritic spine density and morphology were analyzed in Golgi-Cox-impregnated pyramidal neurons located in layer V of the cortex (mCTX+sCTX) at P11 (n= 131 cells WT; 84 cells SMA. Figure 4A, 4B). Quantitative analysis revealed a significant reduction in dendritic spine density in SMA layer V neurons compared to WT controls (WT 3.57 ± 0.06 spines/10µm; SMA 3.13 ± 0.07 spines/10 µm; t-test, p< 0.0001****; Figure 4C). In contrast, the overall spine density along the apical dendrite did not reach statistical significance (WT 5.18 ± 0.12 spines/20µm; SMA 4.79 ± 0.17 spines/20µm; t-test, p=0.08 ns. N= 124 cells WT; 47 cells SMA) (Figure 4D). However, a significant increase in the distance between the soma and the first detectable spine was observed in SMA neurons along both basal and apical dendrites (basal dendrites; WT 3.52 ± 0.1µm; SMA 8.05 ± 0.33µm; t-test, p<0.0001****; apical dendrites; WT 5.77 ± 0.51µm; SMA 10.37 ± 0.4µm; t-test, p<0.0001****; Figures 4E).

**Figure 4.**
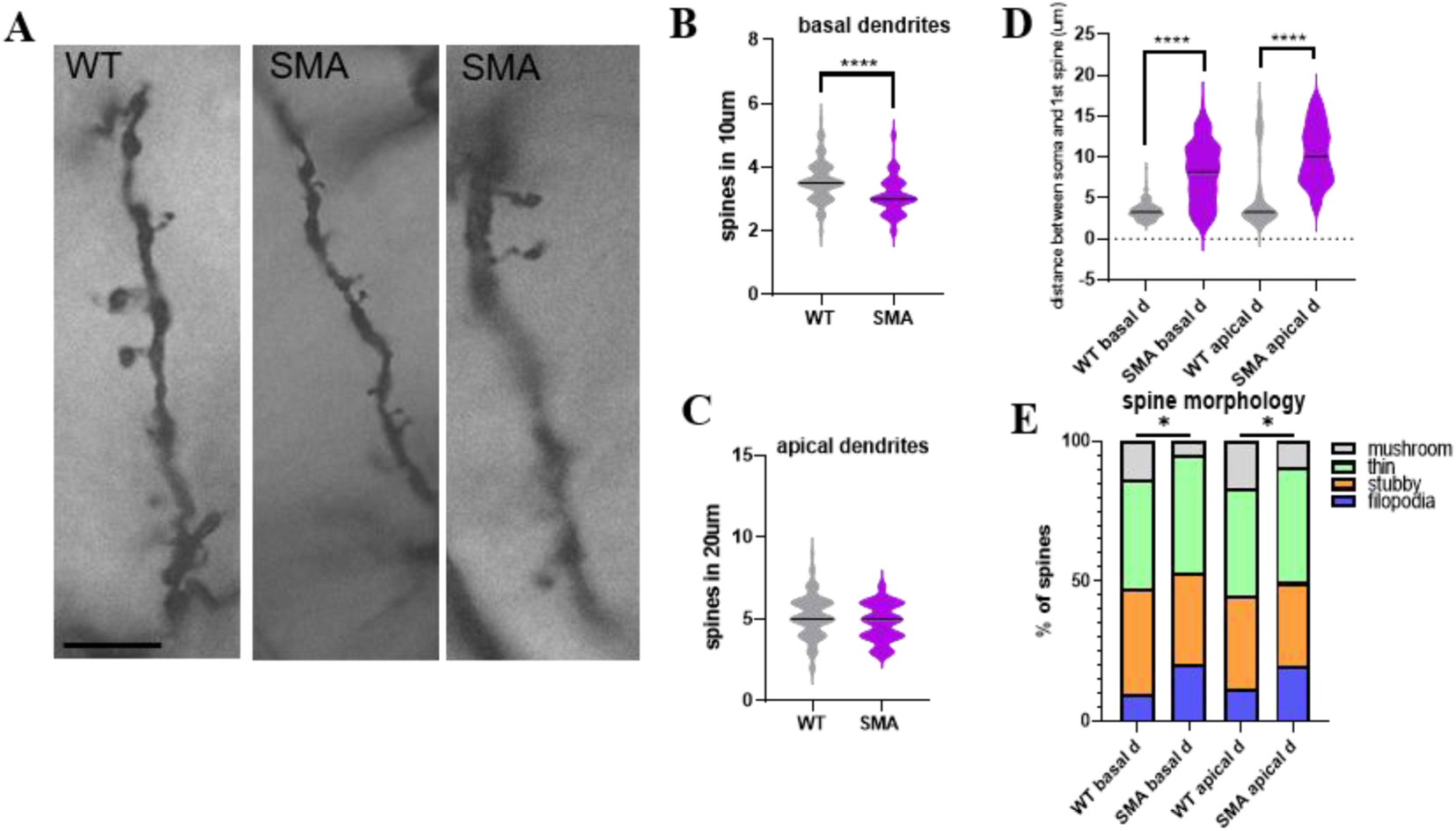
Altered dendritic spine density, distribution, and morphology in layer V cortical neurons of SMA mice at the late symptomatic stage (P11). (A) Representative images from Golgi-Cox-stained layer V pyramidal neurons showing dendritic spine morphology in one WT and two representative SMA neurons at P11. (B) Quantification of dendritic spine density along basal dendrites shows a significant reduction in SMA neurons compared to WT controls. (C) Spine density along apical dendrites does not differ significantly between genotypes. (D) Quantification of the distance between the soma and the first detectable spine reveals a significant increase in SMA neurons in both basal and apical dendrites. (E) Analysis of dendritic spine morphology shows a shift in spine type maturation in SMA neurons, with a reduction in mature mushroom spines and a relative increase in immature spine forms in both basal and apical dendrites. Data are presented as mean ± SEM. Scale bar: 10 µm.

In addition to the reduction in spine density, SMA neurons displayed alterations in spine morphology distribution (Figure 4F): classification of dendritic spines according to established morphological criteria showed a shift in the relative distribution of spine types in both basal and apical dendrites. In particular, mature mushroom spines were reduced, whereas immature spine forms such as filopodia were relatively increased in SMA neurons (basal dendrites; WT, filopodia: 9%, stubby: 38%, thin: 39%, mushroom: 14%; SMA, filopodia: 20%, stubby: 33%, thin: 42%, mushroom: 5%; two-ways ANOVA, p= 0.038*; apical dendrites; WT, filopodia: 11%, stubby: 34%, thin: 38%, mushroom:17%; SMA, filopodia: 20%, stubby: 30%, thin: 41%, mushroom: 9%; two-ways ANOVA, p= 0.03*). This pattern is consistent with impaired synaptic maturation and reduced stability of excitatory synapses.

### Cortical projection neuron population density is preserved in SMA brain at early symptomatic stage

Given the alterations observed in cortical projection neurons at the late symptomatic stage, we asked whether similar changes could already be detected at earlier stages of the disease, by performing quantitative analyses of layer V neurons in SMNΔ7 mice at P5. The analysis of Nissl-stained large pyramidal neurons in layer V revealed no significant differences in neuronal density between SMA mice and WT littermates (WT 1596 ± 36.84 cells/mm^2^; SMA 1582 ± 41.34 cells/mm^2^; t-test, p>0.05ns; n= 6 WT; 4 SMA. Figure 5A-A’). This was observed both in mCTX and sCTX, indicating that total projection neuron density is preserved at this early stage (Figure 5A’’; p>0.05ns).

**Figure 5.**
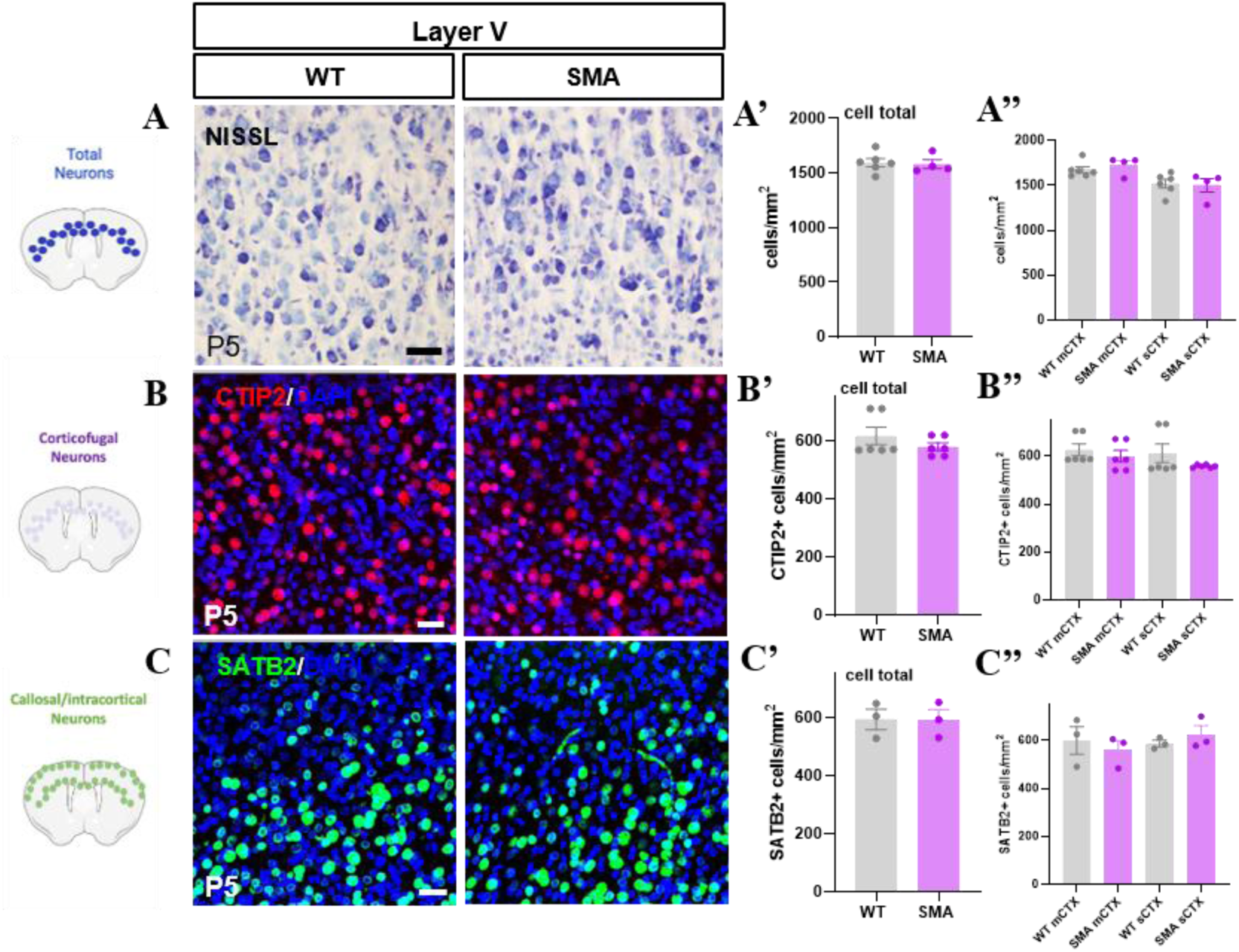
Preservation of cortical projection neuron populations at the early symptomatic stage (P5). (A) Representative Nissl-stained sections of motor cortex from WT and SMA mice at P5. (A’-A’’) Quantification of large pyramidal neurons in layer V shows no significant differences in neuronal density between genotypes, in total and in both motor (mCTX) and somatosensory (sCTX) areas. (B-B’’) Representative immunofluorescence images and quantification of CTIP2-positive corticofugal neurons in layer V reveal comparable densities between WT and SMA mice at P5. (C–C’’) Representative immunofluorescence images and quantification of SATB2-positive callosal projection neurons show no significant differences between genotypes. Data are presented as mean ± SEM (ns, p > 0.05). Scale bars: 50 µm.

To determine whether specific projection neuron subtypes might exhibit early vulnerability, we next quantified corticofugal and callosal/intracortical neuron populations using subtype-specific markers. The density of CTIP2-positive neurons, identifying corticofugal projection neurons, was similar in WT and SMA mice in both cortical regions analyzed (WT 616.1 ± 29.84 cells/mm^2^; SMA 579.4 ± 13.38 cells/mm^2^; t-test, p>0.05ns; n=6 WT; 6 SMA. Figures 5B-B’’). Likewise, SATB2-positive neurons, corresponding to intracortical projection neurons, showed no significant differences between genotypes (WT 593.8 ± 35.10 cells/mm^2^; SMA 592.1 ± 35.16 cells/mm^2^; t-test, p>0.05ns; n= 3 WT; 3 SMA. Figures 5C-C’’). Overall, quantitative analyses confirmed that the density of cortical projection neuron populations was unchanged at P5.

In agreement with these findings, cleaved caspase-3 analysis in CTIP2- and SATB2-positive neuronal populations did not reveal significant differences between SMA and WT mice at P5 (p>0.05ns; Supplementary Fig. 3A, 3B). Despite the relatively small sample size (n = 3) and the high variability observed, these results indicate that clear apoptotic activation is not yet detectable at this early stage, although initial indications of divergence begin to emerge during disease progression.

### SMA corticospinal neurons exhibit early morphological alterations compared to WT

Although the overall density of cortical projection neurons was preserved at P5, morphological analysis of DiI-traced corticospinal and callosal neurons revealed early detectable structural alterations in SMA mice, particularly affecting corticospinal neurons (Fig. 6A-B).

**Figure 6.**
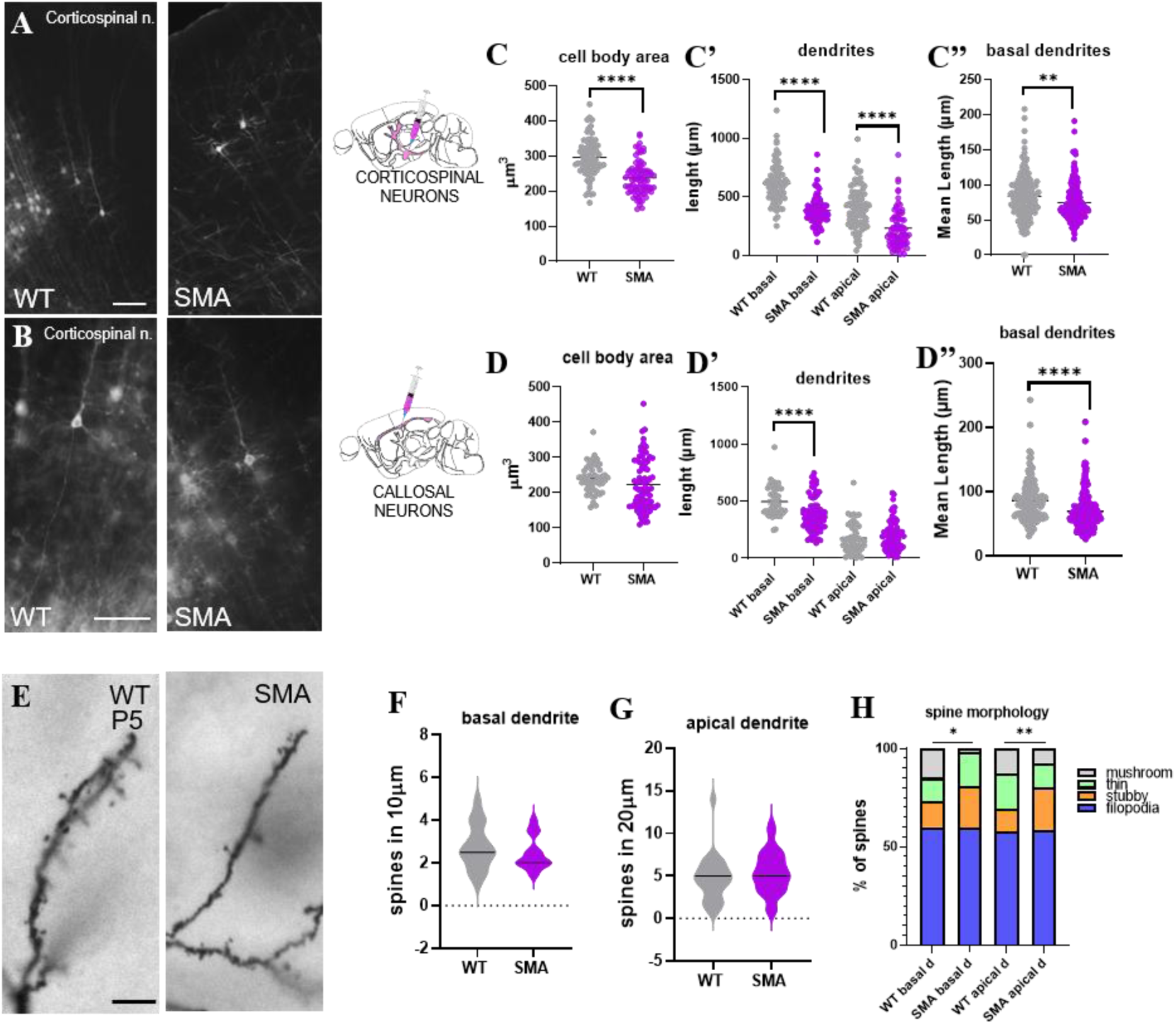
SMA corticospinal neurons exhibit morphological alterations in branching and dendritic spine maturation already in the early phase of the disease. (A-B) Representative fluorescence images of DiI-labeled corticospinal neurons in WT and SMA P5 cortex following DiI injection into the cerebral peduncle. (C-C’’) Quantification of neuronal morphology shows early alterations in SMA corticospinal neurons, including reduced soma size (C) and simplified dendritic architecture and length (C’, C’’) compared to WT controls. (D–D’’) Quantification of neuronal morphology shows a significative reduction in basal dendrite length in SMA callosal neurons compared to WT cells. (E) Representative high-magnification images of basal dendritic segments of Golgi-Cox stained WT and SMA projection neurons showing spines. Quantification of dendritic spine density along basal (F) or apical (G) dendrites shows no significant differences between SMA projection neurons and WT controls. (H) Analysis of dendritic spine morphology shows a shift in spine type maturation in SMA neurons also at early time points, with a significant reduction in mature spines and a relative increase in immature spines, in both basal and apical dendrites. Data are presented as mean ± SEM. Scale bars: 50 µm (A, B); 10 µm (E).

Indeed, quantitative morphometric analysis showed that corticospinal neurons in the SMA cortex exhibited a reduction in soma size compared to WT neurons (n= 79 cells WT; 78 cells SMA. Table 2; Figure 6C), indicating early cellular alterations. In addition, both basal and apical dendrites appeared shorter and less branched, resulting in a decrease in overall dendritic complexity (Table 2; Figures 6C’-C’’ and Supplementary Fig. 3C-C’). These differences were evident despite the absence of significant neuronal loss at this stage.

**Table 2.** Morphological parameters of DiI-labelled corticospinal and corticothalamic neurons at P5.

| Morphological Parameter | Wild Type<br>(Mean ± SEM) | SMA<br>(Mean ± SEM) | p-value<br>(t-test) |
| --- | --- | --- | --- |
| ***Corticospinal neurons (Fig. 6 C- C''')*** |  |  |  |
| Soma Area ($\mu\text{m}^2$) | 262.5 ± 4.57 | 223.8 ± 4.49 | p<0.0001**** |
| Total basal dendrite length ($\mu\text{m}$) | 464.0 ± 15.75 | 364.0 ± 8.1 | p<0.0001**** |
| Basal dendrites mean length ($\mu\text{m}$) | 83.77 ± 2.41 | 74.94 ± 1.67 | p=0.002** |
| Apical dendrite length ($\mu\text{m}$) | 308.4 ± 14.44 | 231.8 ± 10.32 | p<0.0001**** |
| ***Callosally-projecting neurons (Fig. 6 D- D''')*** |  |  |  |
| Soma Area ($\mu\text{m}^2$) | 239.6 ± 5.74 | 223.7 ± 8.074 | p=0.161 ns |
| Total basal dendrite length ($\mu\text{m}$) | 456.5 ± 13.40 | 333.1 ± 9.88 | p<0.0001**** |
| Basal dendrites mean length ($\mu\text{m}$) | 86.58 ± 2.82 | 69.84 ± 2.06 | p<0.0001**** |
| Apical dendrite length ($\mu\text{m}$) | 175 ± 19.05 | 198 ± 13.93 | p=0.322 ns |

In contrast, morphometric analysis of callosal projection neurons at P5 revealed largely preserved structural features between SMA and WT animals (n= 49 cells WT; 82 cells SMA). No significant differences (p> 0.05) were observed in soma size and apical dendrite length (Table 2; Figures 6D-D’), indicating that the overall morphology of this neuronal population is maintained at this stage. Notably, the only parameter showing a significant alteration was the length of basal dendrites, which was reduced in SMA neurons compared to WT controls (Table 2; Figure 6D-6D’’). In accordance with these findings, branching complexity was largely comparable between genotypes for this projection neuron subtype, except for basal dendrites, which showed reduced complexity in SMA neurons (WT 1532 ± 125.6 complexity index; SMA 1034 ± 75.21 complexity index; t-test, p=0.0004***; Supplementary Fig. 3D-D’).

To further assess early synaptic changes in cortical projection neurons, dendritic spine density and morphology were analyzed in Golgi-Cox-impregnated layer V pyramidal neurons also at P5 (Figure 6E). Quantitative analysis revealed no significant differences (p> 0.05) between SMA and WT neurons in basal and apical dendritic spine number (n= 22 cells WT; 25 cells SMA. Figures 6F, 6G), nor in the distance between the soma and the first detectable spine (not shown), suggesting that both synaptic density and proximal spine distribution are preserved at this early stage.

However, classification of spine morphology showed an early shift in the distribution of spine types in SMA neurons. In particular, already in the early phase of the disease, SMA pyramidal neurons exhibited a relative increase in immature spine morphologies, especially stubby spines, accompanied by a reduction in more mature spine types (mushroom type) (basal dendrites; WT, filopodia: 60%, stubby: 13.3%, thin: 11.7%, mushroom:15%; SMA, filopodia: 60%, stubby: 21.4%, thin: 17%, mushroom: 1.6%; two-ways ANOVA, p= 0.013*; apical dendrites; WT, filopodia: 58%, stubby: 11.5%, thin: 18%, mushroom:12.5%; SMA, filopodia: 59%, stubby: 21%, thin: 12.5%, mushroom: 7.5%; two-ways ANOVA, p= 0.0069**; Figure 6H). This change was detectable despite the absence of significant differences in total spine number. Overall, these findings reveal early alterations in spine maturation in SMA cortical neurons despite preserved spine density and neuronal survival at P5.

## Discussion

SMA has usually considered a lower motor neuron disease, and for decades clinical and experimental research - including our previous work (Schellino et al., 2018; Boido et al., 2023) - has primarily focused on lower motor neuron degeneration as its central neurological hallmark. However, accumulating evidence has challenged this view, indicating that SMA should be more appropriately regarded as a multisystem disorder affecting multiple organs and cell types beyond the motor unit (Hamilton and Gillingwater, 2013; Yeo and Darras, 2020). Human postmortem and neuroimaging studies have revealed abnormalities across cortical, thalamic, brainstem, cerebellar and other supraspinal regions, particularly in individuals with lower *SMN2* copy number (Harding et al., 2015; Mendonça et al., 2019; Mugisha et al., 2023; Groulx-Boivin et al., 2024). Consistently, supraspinal structural and functional alterations have been reported in severe SMA mouse models (Tharaneetharan et al., 2021; Cottam et al., 2022; Gerstner et al., 2026).

Together, these macrostructural and circuit-level observations highlight the need to identify the underlying cellular correlates of brain involvement in SMA. In this regard, cortical projection neurons are of particular interest, given previous observations of Betz cell abnormalities in human SMA and selective loss of layer V pyramidal neurons in SMNΔ7 mice (Araki et al., 2003; d’Errico et al., 2013), together with more recent evidence of altered cortical inhibitory circuitry in motor and somatosensory cortices (Menduti et al., 2026).

Here we show that cortical projection neuron alterations in the SMNΔ7 mouse model are subtype- and stage-dependent and, importantly, that structural and synaptic abnormalities emerge before overt cortical neuronal loss.

At the late symptomatic stage (P11), corticospinal and infragranular callosally-projecting neurons were significantly reduced and showed increased CASP3 activation, whereas corticothalamic neurons were numerically preserved. Surviving corticospinal and callosally-projecting neurons also exhibited marked morphological abnormalities, including reduced soma size, shortened dendrites, and simplified dendritic arborization, while corticothalamic neurons showed only minor alterations, essentially limited to apical dendrite length. These findings extend previous evidence of layer V neuronal loss in SMA (d’Errico et al., 2013) by revealing marked heterogeneity in the susceptibility of cortical projection neuron populations to constitutive SMN deficiency.

Interestingly, this selectivity extends within projection neurons classes: indeed, SATB2-positive infragranular projection neurons were significantly reduced at P11, whereas their supragranular counterparts remained preserved. Thus, even neuronal populations sharing a major target may differ substantially in their response to SMN deficiency. Such differential vulnerability may reflect cell-intrinsic molecular and developmental programs, transcriptional identity, connectivity patterns, metabolic demands, or stress-response capacity (Saxena and Caroni, 2011; Fu et al., 2018; Limone et al., 2024; Cheron et al., 2026; Gautier et al., 2026). Cytoskeletal properties may also be determinant, given the established involvement of SMN in cytoskeletal regulation, axonal transport, and neurite growth (Shi et al., 2024), and laminar position and connectivity may also influence the selective vulnerability of specific cortical projection neuron populations, as shown in other neurodegenerative conditions such as Huntington’s disease (Pressl et al., 2023), as well as their dependence on SMN deficiency. Indeed, given that distinct cortical projection neuron populations are generated and specified at different stages of corticogenesis, they may experience distinct developmental windows of sensitivity to reduced SMN levels, potentially affecting their generation, positioning, or subsequent maturation. Similar links between early synaptic/network dysfunction, altered neurite growth and branching, and subsequent neuronal vulnerability have been described in ALS patient-derived motor neurons (Kollstrøm et al., 2025).

Importantly, neuronal degeneration at P11 was accompanied by alterations in the excitatory synaptic milieu. Reduced vGlut1 immunoreactivity and fewer vGlut1-positive puncta were detected in layer V, and notably, also in supragranular layers despite preservation of the SATB2-positive population, indicating that excitatory abnormalities are not restricted to layers undergoing detectable projection neuron loss. Because vGlut1-positive terminals may originate from multiple cortical and extracortical glutamatergic populations, their source cannot be established from the present analysis; nevertheless, these findings point to a broader alteration in the organization or maintenance of excitatory cortical connectivity, consistent with developmentally evolving functional abnormalities previously reported in SMA motor cortical networks (Tharaneetharan et al., 2020).

Dendritic spine abnormalities further support altered synaptic organization. At P11, SMA layer V neurons displayed reduced basal spine density together with a shift toward immature spine morphologies (fewer mature mushroom spines and a relative enrichment of immature forms), and increased soma-to-first-spine distance. Notably, in apical dendrites, where overall spine density was not significantly reduced, the extended proximal spine-free segment suggests an altered spatial organization of synaptic contacts, potentially reflecting preferential loss of proximal spines or their redistribution along the dendritic shaft. Together with the reduced dendritic arborization and branching complexity observed in layer V neurons, these findings indicate substantial remodeling of the receptive and synaptic architecture of surviving SMA cortical neurons, consistent with a structurally and functionally vulnerable phenotype. The persistence of immature spine morphologies is particularly relevant considering emerging evidence of altered glutamatergic homeostasis in SMA. While our data points to an early disruption of excitatory synaptic inputs, a concomitant contribution of altered glutamatergic signaling and metabolism cannot be excluded. Indeed, SMN deficiency has been associated with reduced glutamate levels, impaired neuron-astrocyte metabolic coupling and hypoglutamatergic neurotransmission (Hassan et al., 2025), suggesting that altered glutamatergic signaling may contribute to the synaptic phenotype observed here. Notably, local glutamate receptor activation promotes the maturation of dendritic protrusions toward mature spine morphologies (Mattison et al., 2014), raising the possibility that reduced glutamatergic signaling and impaired spine maturation are therefore mechanistically interconnected in SMA, although our data cannot establish the directionality of this relationship. These excitatory abnormalities likely occur within a wider network disturbance, as cortical GABAergic dysfunction (Menduti et al., 2026) and excitation/inhibition imbalance (Fletcher et al., 2025) have also been reported in SMA. Moreover, motor cortical network abnormalities evolve during development, supporting a potential role for early circuit dysfunction rather than an exclusively secondary consequence of motor neuron degeneration (Tharaneetharan et al., 2020). This concept is consistent with evidence from other neurodegenerative disorders, including amyotrophic lateral sclerosis (ALS), in which early synaptic alterations and circuit dysfunction may contribute to selective neuronal vulnerability (Arnold et al., 2024; Salzinger et al., 2024; Kollstrøm et al., 2025; Tessitore et al., 2026).

A central finding of our study is that cortical abnormalities emerge progressively before overt neuronal loss and are not restricted to late stages of the disease. At P5, projection neuron density was preserved, and CASP3 activation did not significantly differ between genotypes, although higher positivity was already apparent in CTIP2- and SATB2-positive populations, supporting the idea of early impairment preceding degeneration. Despite preserved cell numbers, corticospinal neurons already exhibited smaller somata, shorter dendrites and axons, and reduced dendritic complexity, whereas callosal neurons were comparatively preserved, showing milder alterations mainly involving basal dendrites. Notably, the soma size of SMA corticospinal neurons failed to increase between P5 and P11, suggesting a failure of normal neuronal development, and indicating that these neurons may not reach full structural maturity in the absence of SMN. Thus, the most vulnerable populations of cortical projection neurons already display signs of altered postnatal maturation before neuronal loss becomes detectable.

This differential response suggests a temporal hierarchy of vulnerability among cortical projection neurons, in which corticospinal neurons exhibit the earliest structural abnormalities, callosal neurons become more extensively affected during progression, and corticothalamic neurons remain comparatively resistant even at P11. The analysis of dendritic spines reinforces this developmental interpretation. Although dendritic spine density is not altered at P5, SMA neurons already exhibit a shift toward immature spine phenotypes that persists during disease progression and precedes the reduction in spine density observed at later stages. Because spine maturation is highly dynamic during postnatal cortical development and essential for circuit formation (Egashira et al., 2026), these findings may reflect impaired developmental maturation rather than simply secondary degeneration of established synapses. Such early synaptic vulnerability preceding neuronal loss has similarly been described across several neurological conditions (Wishart et al., 2006; Henstridge et al., 2016; Taoufik et al., 2018), including Parkinson’s disease (Belloso-Iguerategui et al., 2023; Rodriguez-Aller et al., 2025), Alzheimer’s disease (Shankar and Walsh, 2009; Pelucchi et al., 2022) and ALS, in both spinal cord and cortex (Sasaki and Maruyama, 1994; Nagao et al., 1998; Sasaki and Iwata, 1999).

This interpretation is consistent with increasing evidence that SMN deficiency interferes with neurodevelopment. Early defects in neuronal differentiation and network activity have been reported in patient-derived SMA organoids before overt cell death (Grass et al., 2024; Faravelli et al., 2025), while developmental abnormalities have also been identified in SMA mouse brain (Wishart et al., 2010). Cognitive, behavioral, and neurodevelopmental alterations have increasingly been reported clinically, particularly in severe SMA, although their prevalence and direct relationship with SMN deficiency remain debated, because primary abnormalities are difficult to distinguish from consequences of profound early motor impairment (Masson et al., 2021; Gómez-Andrés et al., 2026; Gnazzo et al., 2026).

The temporal relationship between cortical and spinal pathology nevertheless remains an important unsolved question. Overt cortical projection neuron degeneration becomes evident at P11, whereas spinal motor neuron pathology is established earlier in the SMNΔ7 model (Butchbach et al., 2007; Bäumer et al., 2009; d’Errico et al., 2013). Cortical neuronal loss could therefore be temporally downstream of spinal degeneration and be influenced by progressive disruption of interconnected motor circuits. However, morphological and synaptic abnormalities in cortical neurons already present at P5 argue against an exclusively secondary origin. Our findings, therefore, favor a model in which constitutive SMN deficiency establishes an early intrinsic and developmental vulnerability of cortical neurons, while progressive alterations in spinal and supraspinal connectivity may subsequently modulate or exacerbate their degeneration (Figure 7). Primary cellular vulnerability and secondary network-dependent mechanisms need not be mutually exclusive but may interact throughout disease progression.

**Figure 7.**
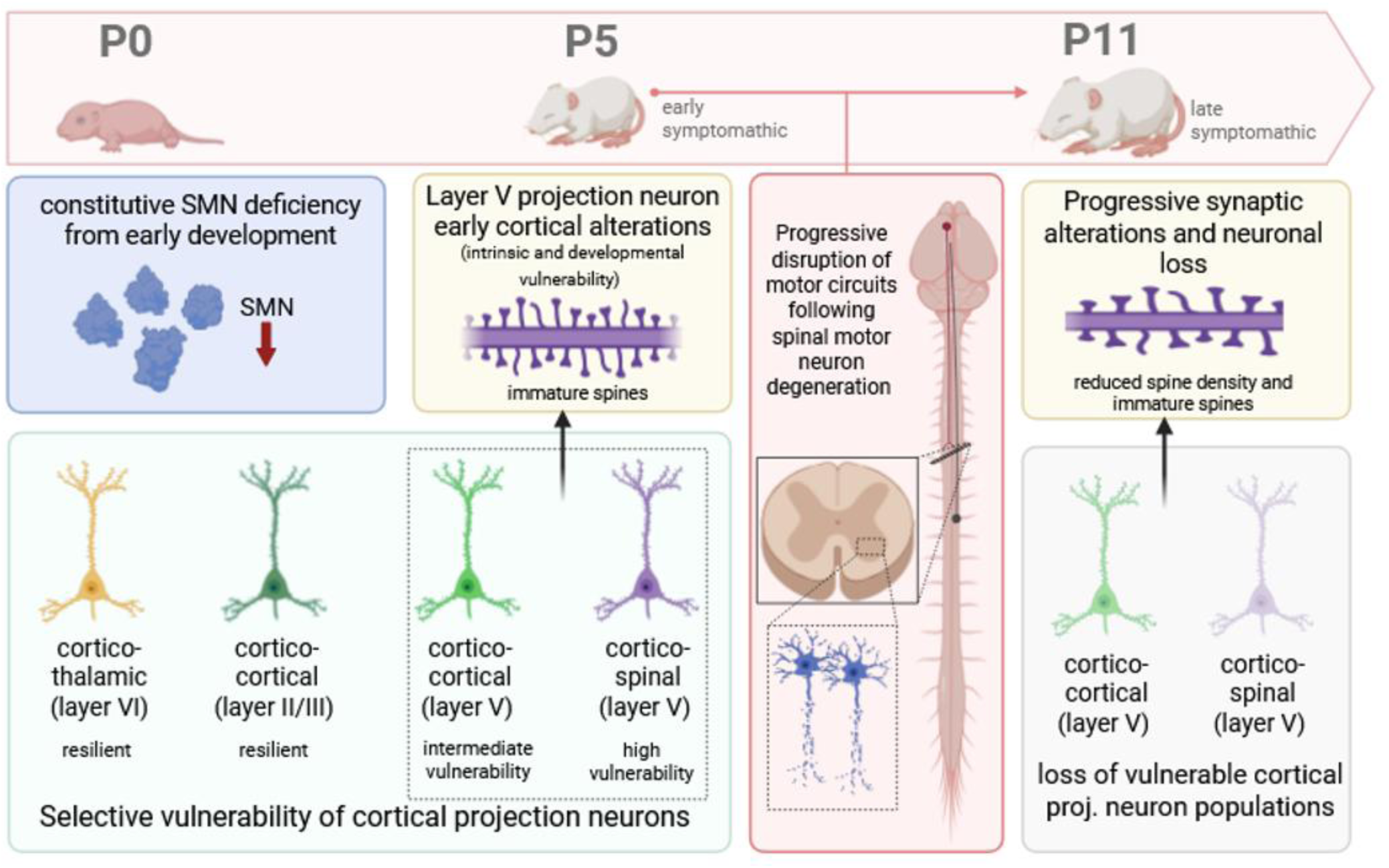
Proposed working model of the temporal progression and selective vulnerability of cortical projection neurons in SMA. Schematic representation of the main findings and proposed mechanisms underlying cortical involvement in the SMNΔ7 mouse model. Constitutive SMN deficiency is associated with early structural and synaptic maturation defects in cortical projection neurons, already detectable at P5 before overt neuronal loss, with corticospinal neurons showing the greatest vulnerability, followed by corticocortical neurons in layer V. During disease progression, the well-established spinal motor neuron degeneration and associated disruption of interconnected motor circuits may additionally modulate cortical pathology. At the late symptomatic stage (P11), early abnormalities are accompanied by reduced spine density and selective loss of vulnerable layer V corticospinal and corticocortical projection neurons, whereas supragranular corticocortical and corticothalamic populations remain comparatively preserved. The proposed model therefore suggests that cortical pathology may arise from the interplay between early intrinsic and developmental vulnerability to SMN deficiency and subsequent network-dependent alterations, rather than representing exclusively a secondary consequence of spinal motor neuron degeneration. Created with BioRender.com.

Lastly, several limitations should be considered. CTIP2 and SATB2 transcription factors provide useful molecular proxies for major cortical projection neuron populations but their expression partially overlaps across subclasses and developmental stages; therefore, marker-based quantification approximates rather than unequivocally defines projection identity. Projection-specific DiI tracing provided complementary anatomical identification for the morphometric analyses, but technical constraints related to protocol compatibility precluded its combination with marker-based cell quantification. In addition, this primarily morphological and histological study cannot directly establish functional consequences or causal relationships among early maturation defects, circuit alterations, and neuronal loss. Finally, because our postnatal analyses cannot determine whether the observed abnormalities already originate during embryonic development. These limitations define important future directions. In particular, projection-defined single-cell or single-nucleus transcriptomic and other omics-based approaches could identify molecular programs associated with vulnerability or resilience among cortical projection neuron subtypes, including developmental and corticogenic pathways differentially sensitive to reduced SMN levels. Integrating these molecular profiles with the temporal and morphological phenotypes identified here may help clarify how altered developmental trajectories contribute to the selective vulnerability of specific cortical populations.

The temporal component also has therapeutic implications. Current SMN-restoring approaches represent a major advance in SMA treatment, and early SMN restoration preserves spinal motor neuron survival and function in experimental models (Aslesh and Yokota, 2022; Berciano et al., 2020; Comley et al., 2022). Early antisense treatment can also rescue developmental abnormalities in patient-derived SMA spinal cord organoids (Faravelli et al., 2025), supporting the reversibility of at least some early CNS phenotypes. Determining whether early SMN restoration can similarly normalize cortical projection neuron growth, dendritic complexity, spine maturation, and ultimately neuronal survival in vivo will be important for defining the reversibility and therapeutic window of supraspinal alterations.

Overall, our findings expand the pathophysiological spectrum of SMA beyond lower motor neuron degeneration, demonstrating selective cortical projection neuron vulnerability to constitutive SMN deficiency. Importantly, early impairments in neuronal and synaptic maturation precede overt degeneration, supporting a model in which developmental and intrinsic neuronal susceptibility interacts with progressive network dysfunction to shape supraspinal pathology and potentially contribute to disease progression. Defining the molecular determinants and reversibility of these early cortical abnormalities may ultimately help extend therapeutic strategies beyond spinal motor neuron preservation to the broader neural circuits affected by SMN deficiency.

## Author Contributions

Conceptualization: RS, MB, AV; Methodology: all authors; Investigation and data curation: RS, FLR, GI, AC; Writing - Original Draft: RS, FLR, GI; Writing - Review & Editing: RS, FLR, MB, AV; Supervision: RS, MB, AV; Funding Acquisition: MB, AV.

