## Supplementary Figures for "Cortical projection neurons exhibit early maturation defects and selective degeneration in a mouse model of spinal muscular atrophy"

### Supplementary Figure 1

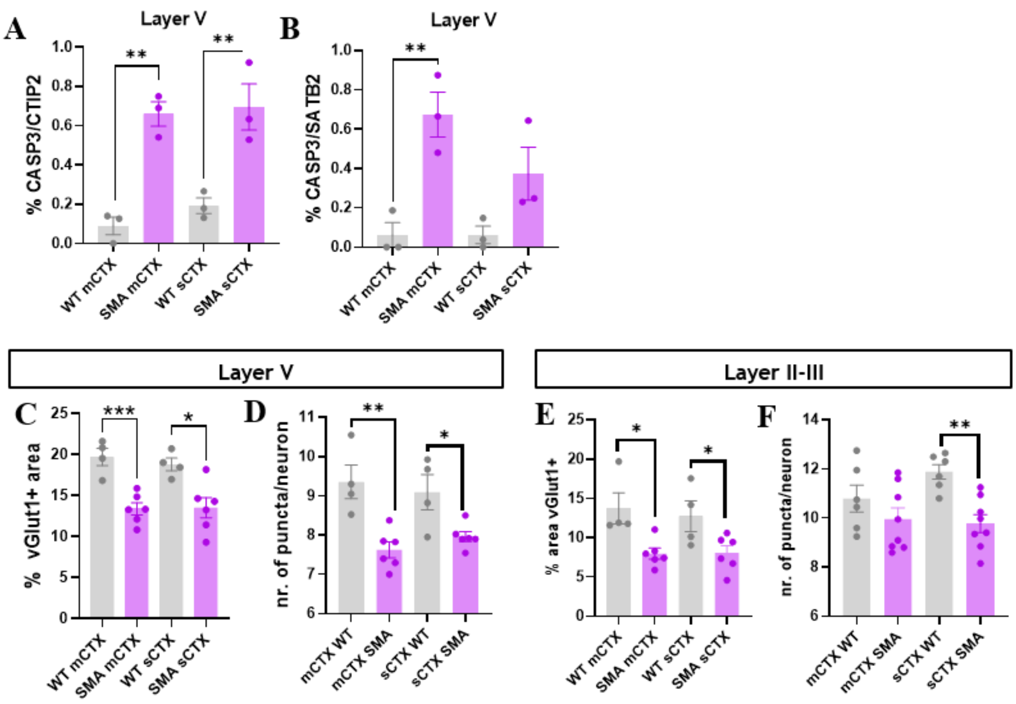


**Supplementary Figure 1.** (A) Quantification of CASP3-positive CTIP2+ corticofugal neurons in motor (mCTX) and somatosensory (sCTX) cortices of P11 WT and SMA mice. (B) Quantification of CASP3-positive SATB2+ callosal neurons in motor and somatosensory areas of WT and SMA littermates at P11. (C) Quantification of vGlut1-positive signal intensity in layer V of motor and somatosensory cortices, revealing a reduction in SMA mice compared to WT.(D) Quantification of vGLUT1-positive puncta per NeuN-positive neuron in layer V at P11, showing decreased excitatory synaptic input in SMA cortex across both regions.(E) Quantification of vGlut1-positive signal intensity in supragranular layers (II–III) of motor and somatosensory cortices in P11 WT and SMA mice.(F) Quantification of vGlut1-positive puncta in supragranular layers, showing reduced puncta in SMA mice, with a more pronounced effect in somatosensory areas. Data are presented as mean ± SEM. Statistical analysis was performed using unpaired Student's t-test (*p < 0.05, **p < 0.01, ***p < 0.001).

### Supplementary Figure 2

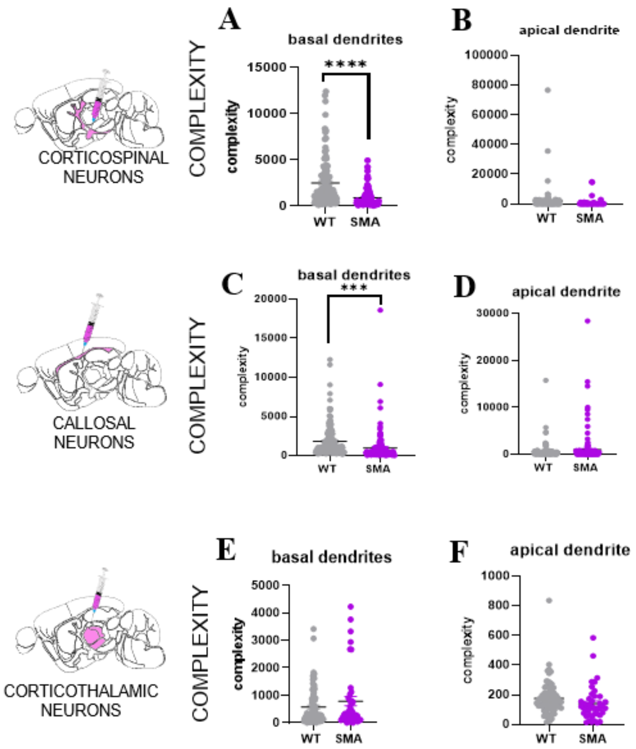


**Supplementary Figure 2.** (A-B) Neurolucida Explorer-derived analyses of dendritic complexity (basal and apical dendrites) in corticospinal neurons show a marked decrease in complexity index in SMA basal dendrites. (C-D) Neurolucida Explorer-derived analyses of dendritic complexity (basal and apical dendrites) in callosal neurons also show a significative decrease in complexity index in SMA basal dendrites. (E-F) Neurolucida Explorer-derived analyses of dendritic complexity (basal and apical dendrites) in corticothalamic neurons show no differences between WT and SMA. Data are presented as mean ± SEM. Statistical analysis was performed using unpaired Student's t-test (*p < 0.05, **p < 0.01, ***p < 0.001).

### Supplementary Figure 3

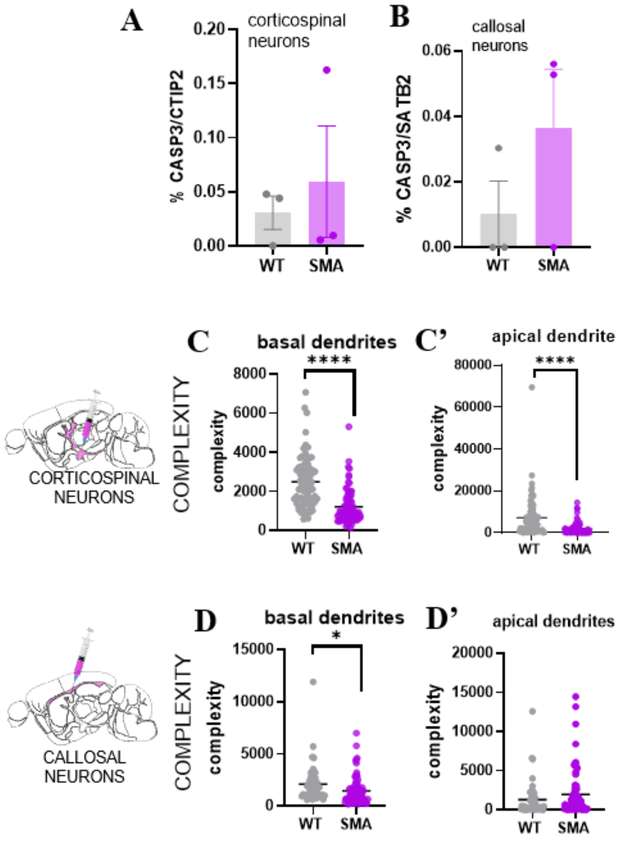


**Supplementary Figure 3.** (A-B) Quantification of CASP3-positive cells in corticospinal (CTIP2$^{+}$, A) and callosal (SATB2$^{+}$, B) projection neurons at P5 shows no significant differences between WT and SMA mice, although a trend toward increased apoptotic signaling is observed in SMA neurons. (C-D) Neurolucida Explorer–based analysis of neurite complexity in DiI-labeled corticospinal (C-C') and callosal (D-D') neurons at P5 reveals early structural alterations in SMA neurons, characterized by reduced dendritic complexity compared to WT controls, particularly in corticospinal neurons. Data are presented as mean ± SEM. Statistical analysis was performed using unpaired Student's t-test (*p < 0.05, **p < 0.01, ***p < 0.001).
